# Platelet Derived Growth Factor Receptor β (PDGFRβ) is a Host Receptor for the human malaria parasite adhesin TRAP

**DOI:** 10.1101/2020.05.12.091488

**Authors:** Ryan W. J. Steel, Vladimir Vigdorovich, Nicholas Dambrauskas, Brandon K. Wilder, Silvia A. Arredondo, Debashree Goswami, Sudhir Kumar, Sara Carbonetti, Kristian E. Swearingen, Thao Nguyen, Will Betz, Nelly Camargo, Bridget S. Fisher, Jo Soden, Helen Thomas, Jim Freeth, Robert L. Moritz, D. Noah Sather, Stefan H.I. Kappe

**Author notes:** Equal contribution (Ryan W. J. Steel, Vladimir Vigdorovich). Infectious Disease and Immune Defence Division, The Walter and Eliza Hall Institute of Medical Research, Parkville 3052, VIC, Australia. Vaccine and Gene Therapy Institute, Oregon Health & Science University, Beaverton, OR 97006. To whom correspondence should be addressed: Stefan Kappe;, D. Noah Sather.

## Abstract

Following their inoculation by the bite of an infected *Anopheles* mosquito, the malaria parasite sporozoite forms travel from the bite site in the skin into the bloodstream, which transports them to the liver. The thrombospondin-related anonymous protein (TRAP) is a type 1 transmembrane protein that is released from secretory organelles and relocalized on the sporozoite plasma membrane. TRAP is required for sporozoite motility and host infection, and its extracellular portion contains adhesive domains that are predicted to engage host receptors. Here, we identified the human platelet-derived growth factor receptor β (hPDGFRβ) as one such protein receptor. Deletion mutants showed that the von Willebrand factor type A and thrombospondin repeat domains of TRAP are both required for optimal binding to hPDGFRβ-expressing cells. We also demonstrate that this interaction is conserved in the human-infective parasite *Plasmodium vivax*, but not the rodent-infective parasite *Plasmodium yoelii*. We observed expression of hPDGFRβ mainly in cells associated with the vasculature suggesting that TRAP:hPDGFRβ interaction may play a role in the recognition of blood vessels by invading sporozoites.

## Introduction

There were more than 400,000 deaths and 228 million estimated new clinical cases of malaria in 2019, the vast majority of which were caused by *Plasmodium falciparum* and *P. vivax* (1). Malaria infection begins when ≤100 sporozoites are deposited into the skin by the bite of a female *Anopheles* mosquito (2). While in skin, sporozoites search for blood vessels using unknown molecular interactions, and then invade the vasculature, through which they are transported to the liver. Here, sporozoites extravasate by traversing liver sinusoidal cells and Kupffer cells, and subsequently enter the liver parenchyma (3, 4). In the liver, each sporozoite traverses several hepatocytes before ultimately selecting one suitable for infection, within which it transforms and grows as liver-stage parasite, ultimately differentiating into tens of thousands of exoerythrocytic merozoites. Upon egress from the liver, these merozoites go on to infect red blood cells. The ensuing erythrocytic infection cycle rapidly increases the parasite load and is responsible for all clinical malaria symptoms, as well as for giving rise to mosquito infectious stages ready for uptake by a new mosquito vector (5–9). Thus, the sporozoite stage is considered an attractive target for intervention, since preventing a relatively small number of parasites from reaching the liver promises to prevent both disease and onwards transmission of the parasite.

Parasite adhesion proteins enabling sporozoite motility, tissue traversal and infection are released from a set of apical organelles, the micronemes and rhoptries (10). An important sporozoite micronemal protein called thrombospondin-related anonymous protein (TRAP), is relocalized from the micronemes to the sporozoite surface and mediates motility and infection via interaction of its cytoplasmic domain with the actin/myosin motor (11). The extracellular portion of TRAP contains two adhesive domains, a von Willebrand factor A (VWA)-like domain with a metal ion-dependent adhesion site (MIDAS), and a thrombospondin repeat domain (TSR domain). TSR and VWA domains in other organisms are broadly involved in receptor binding, cell-cell recognition and cell adhesion, suggesting a similar role for TRAP in binding host receptors. The TSR domain of TRAP has been shown to bind to heparan sulfate proteoglycans (HSPGs) on the surface of hepatocytes (12). Proteolytic processing of TRAP at a rhomboid protease site positioned between the repeats and the transmembrane domain (Fig. 1A) has been shown to be important for sporozoite motility (13).

**Figure 1:**
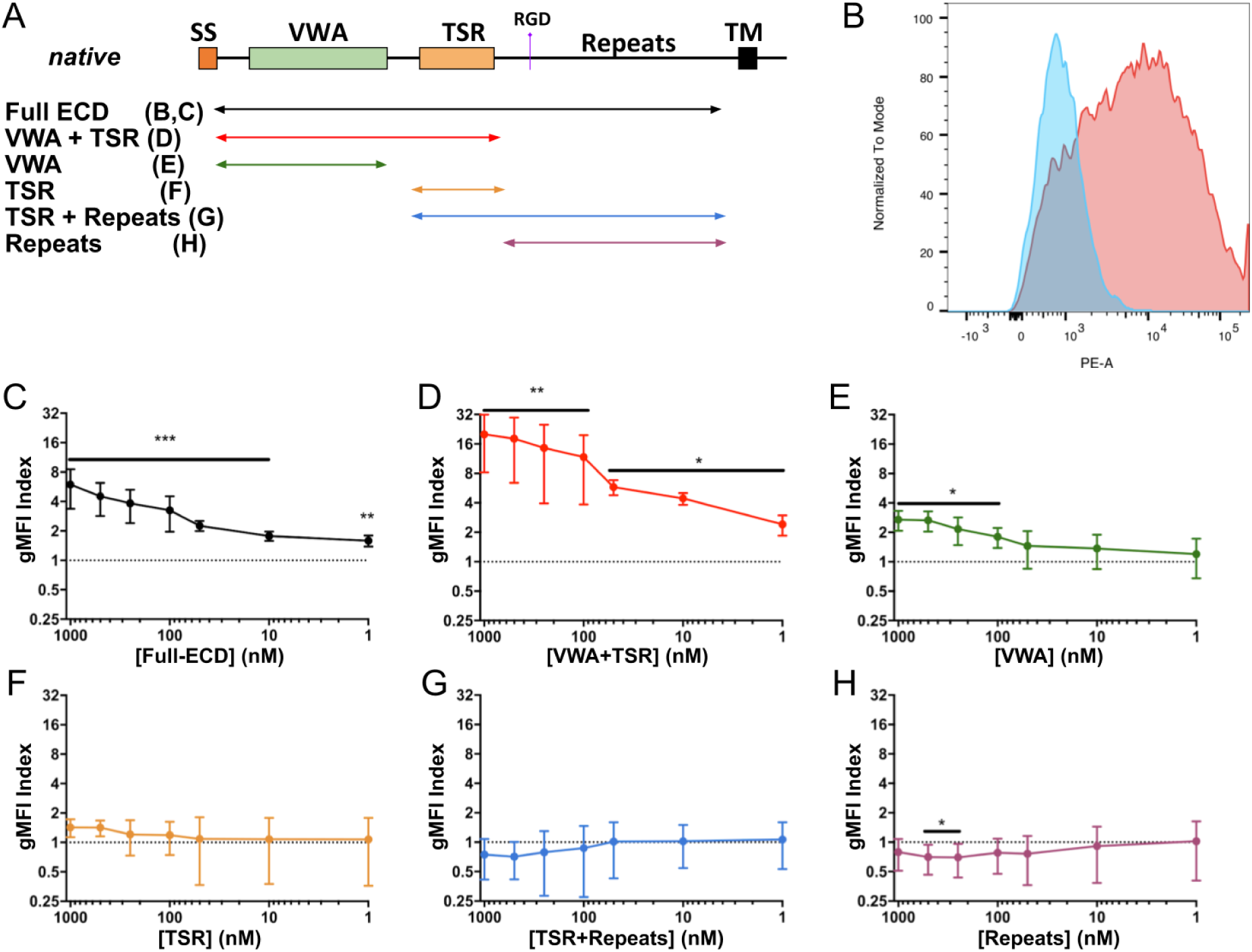
The VWA and TSR domains of PfTRAP act together to bind human PDGFRβ. (A) Overlapping constructs containing various domains of TRAP are represented as a schematic and were similarly used in further experiments. (B) Biotinylated and fluorescently-labeled PfTRAP ectodomain binds preferentially to the HEK293F cells transfected with hPDGFRβ (red) over untransfected cells (blue) within the same culture. Titration experiments (C–H) using increasing dilution of fluorescently labeled fragments of PfTRAP were quantitated as a ratio of binding to transfected versus untransfected cells (gMFI Index) across a 3-log concentration range. Fragments of PfTRAP containing the VWA and TSR domains together bound PDGFRβ strongly across a 3-log range (C, D). While the VWA domain alone showed significant binding to hPDGFRβ at concentrations of 100 nM and above (E), the TSR fragment, the repeat region, or the TSR fragment plus the repeat region did not bind significantly (F–H). Data are the mean ± SD from at least five independent experiments. Analysis by one sample t-test compared to the dashed line (gMFI index expected if no specific binding occurs); *p<0.05, **p<0.01, ***p<0.001.

TRAP localization in sporozoites is a dynamically controlled process. A small amount of TRAP is released apically from micronemes and subsequently translocated rearward on the sporozoite surface during gliding, acting as an essential link between the sporozoite actin/myosin motor and extracellular substrates (13–15). During hepatocyte infection, TRAP becomes largely surface-exposed at the anterior tip of the sporozoite along with other invasion-related proteins, where it is presumably involved in the engagement of host cell receptors (16). This binding to host receptors has been proposed to follow a ‘dual binding system’ model in which the VWA and TSR domains of TRAP coordinate host receptor engagement to allow efficient host infection (17). Mutations in either of these domains reduce sporozoite infectivity while their simultaneous mutation or deletion renders sporozoites non-motile and non-infectious (17, 18). Crystal structures suggest that TRAP undergoes conformational changes when divalent cations bind to its MIDAS site (19).

The critical role of TRAP during host infection makes it an attractive vaccine candidate, although efforts to generate protective antibody responses against TRAP have achieved only limited success to date (20–24). Thus, a better understanding of the specific host-receptor interactions of TRAP during infection would aid ongoing vaccine development by directing rational vaccine design efforts.

Here we expressed the ectodomain of *P. falciparum* TRAP (PfTRAP) and screened for interaction with >4300 individually overexpressed human cell surface receptors. We found that PfTRAP interacts with human platelet-derived growth factor receptor β (hPDGFRβ) and further investigated this interaction using HEK293F cells overexpressing this receptor. Binding of PfTRAP to hPDGFRβ was inhibited by monoclonal antibodies directed against PfTRAP or hPDGFRβ. We further studied what domains of PfTRAP were required for this interaction, and whether the engagement of PDGFRβ by TRAP is conserved in other *Plasmodium* species.

## Results

### Recombinant *P. falciparum* TRAP interacts with human PDGFRβ

To identify host receptors for PfTRAP, we expressed its full ectodomain (Table 1, Fig. 1A) in HEK293F cells and purified the soluble secreted protein (Suppl. Fig. 1B). Using this protein, we undertook a screen of >4300 unique human plasma membrane proteins using the Retrogenix® Cell Microarray platform. In this screen, individual constructs encoding membrane proteins were reverse-transfected into HEK293 cells resulting in an array of populations overexpressing different targets. These fixed arrays were then interrogated using fluorescent complexes containing the PfTRAP ectodomain in search of human receptors that demonstrated reproducible binding to PfTRAP. This screen revealed hPDGFRβ as a PfTRAP-interacting protein (Suppl. Fig. 2). We subsequently verified this interaction using a flow-cytometry-based binding assay, in which HEK293F cells overexpressing hPDGFRβ were stained with fluorescent complexes containing the PfTRAP ectodomain oligomerized around a phycoerythrin-streptavidin core (Fig. 1B).

**Table 1:** Protein constructs.

| **Protein (Database Reference)** | **Domain(s)** | **Amino Acids** |
| --- | --- | --- |
| PfTRAP (PlasmoDB:PF3D7_1335900) | Full Ectodomain | 26–513 |
|  | VWA+TSR | 26–288 |
|  | VWA | 26–244 |
|  | TSR | 241–292 |
|  | TSR + Repeats | 243–513 |
|  | Repeats | 291–513 |
| PvTRAP (*PlasmoDB:PVX_082735*,<br>*PlasmoDB:PVP01_1218700*)<sup>1</sup> | Full Ectodomain | 25–493 |
| PyTRAP (PlasmoDB:PYYM_1351500) | Full Ectodomain | 23–763 |
|  | VWA+TSR | 23–281 |
| PDGF-BB (GenBank: CAG30424.1) | Ectodomain | 82–190 |
| hPDGFRβ (GenBank: BC032224.1) | Ectodomain | 33–532 |

<sup>1</sup> Protein sequence was modified, as described in the main text.

In order to assess the expression of hPDGFRβ in tissues most relevant for sporozoite infection, we undertook an immunohistochemical analysis of normal human skin and liver using an antibody specific for hPDGFRβ. In the skin, strong hPDGFRβ expression was observed in pericytes that undergird the basement membrane of blood vessels (Fig. 2A). To a lesser degree, hPDGFRβ was also expressed in the basal cell layer of the epidermis. In the liver, more diffuse, low levels of hPDGFRβ were observed in proximity to major vessels but not in association with liver sinusoids (Fig. 2B).

**Figure 2:**
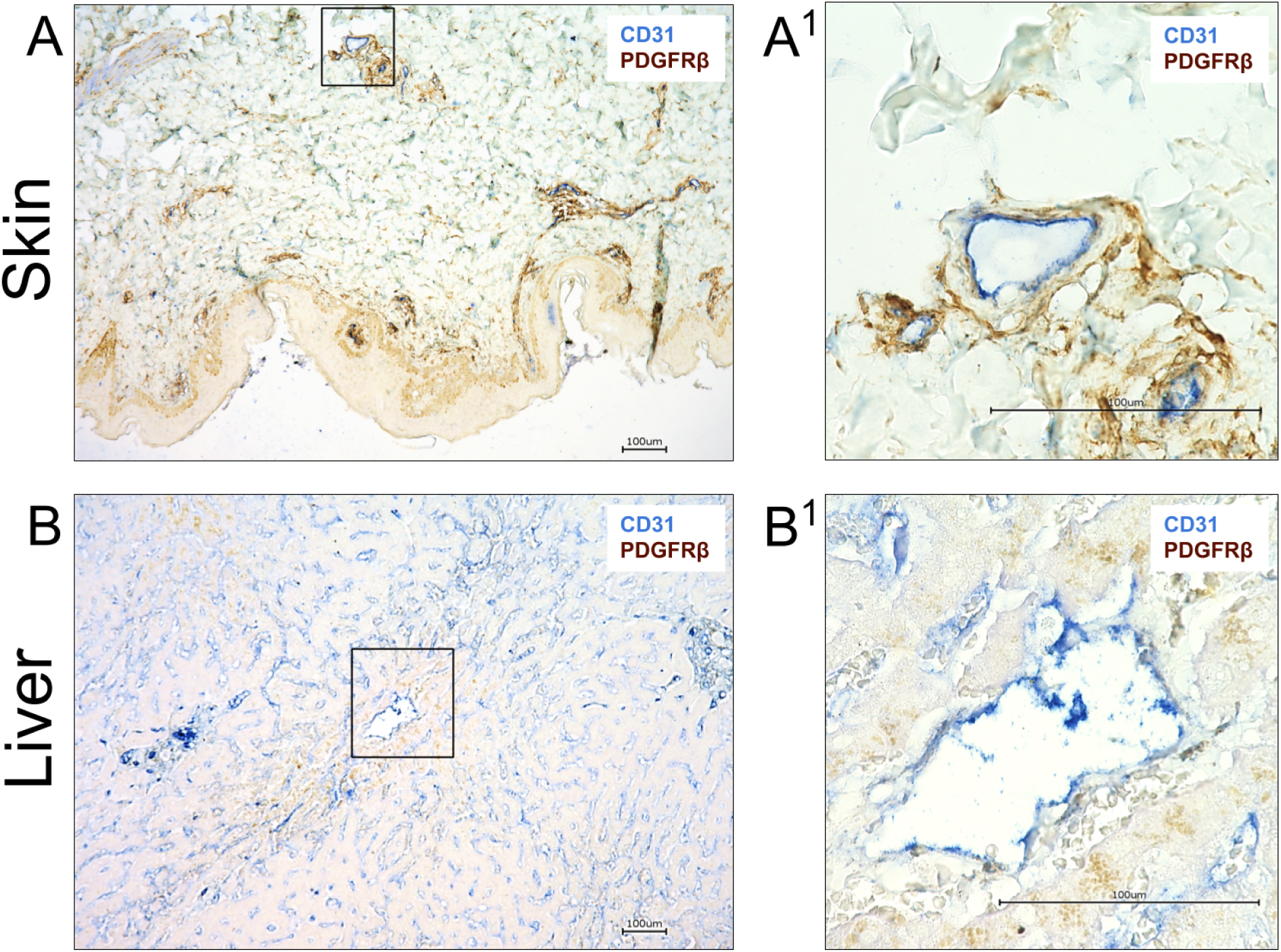
hPDGFRβ is expressed in the basement membrane of vessels in the skin, and around large blood vessels in the liver. Formalin fixed paraffin embedded sections of normal human skin and liver were stained for CD31 (blue) and PDGFRβ (brown). (A) Staining for the endothelial marker CD31 could be seen in the lumen of all skin vasculature irrespective of the size. PDGFRβ was detected surrounding these vascular structures but was not associated with the luminal side of the vessel. Boxed area enlarged in A^1^. (B) CD31 staining was observed in liver sinusoidal endothelial cells (LSEC) all throughout the liver and also in the lumen of major blood vessels. Very low levels of PDGFRβ staining was detected surrounding large blood vessels but was not associated with the luminal surface of the vessel. No PDGFRβ staining was seen around LSECs. Boxed area enlarged in B^1^. Scale bars are 100 µm.

### Optimal engagement of hPDGFRβ by *P. falciparum* TRAP requires the VWA and TSR domains and is independent from the mechanism of binding to αv-containing integrin

We used our flow-cytometry-based binding assay to further investigate the interaction between hPDGFRβ and TRAP (the full gating strategy is shown in Suppl. Fig. 3). In addition to the full-length ectodomain, we expressed and purified five deletion constructs of PfTRAP (Table 1; Suppl. Fig. 1B), as well as the homodimeric form of human platelet derived growth factor B (hPDGF-BB), a known ligand of hPDGFRβ, to be used as a positive control. All purified recombinant proteins were biotinylated and oligomerized using a phycoerythrin-streptavidin conjugate to form a fluorescent staining reagent that was then tested across a 3-log concentration range for binding to HEK293F cells overexpressing hPDGFRβ. Untransfected cells present within each experiment (and trackable due to the lack of GFP signal, which indicated the presence of the fusion hPDGFRβ protein), were used to calculate the geometric mean fluorescence intensity (gMFI) ratio with respect to the cells expressing hPDGFRβ-GFP fusion (referred to as gMFI Index).

Full-length PfTRAP ectodomain bound in a concentration-dependent manner to hPDGFRβ-expressing cells (Fig. 1C). This binding was enhanced (p<0.01; Suppl. Fig. 4) for the VWA+TSR fragment (Fig. 1D). The VWA fragment without the TSR domain bound only modestly (Fig. 1E), while the TSR, alone or in combination with the repeats, or the repeats alone, did not bind to HEK293F cells overexpressing hPDGFRβ (Figs. 1F, G, H).

We next used monoclonal antibodies (mAbs) directed against PfTRAP and hPDGFRβ to further evaluate this interaction. Antibodies that bind the PfTRAP VWA domain reduced the binding of both the full-length ectodomain and the VWA+TSR fragment of PfTRAP by approximately 80% and 90%, respectively, but these mAbs did not affect binding of the known ligand hPDGF-BB to hPDGFRβ (Fig. 3A). We also employed mAb 2C5, which binds to mouse and human PDGFRβ (Suppl. Fig. 5) and blocks activation of these receptors by their natural ligand PDGF-BB *in vitro* and *in vivo* (25). MAb 2C5 significantly reduced the binding of full-length ectodomain and the VWA+TSR fragment, and that of the known ligand, hPDGF-BB, to HEK293F cells overexpressing hPDGFRβ (Fig. 3B). The blocking activity of the antibodies against either partner indicates the specificity of the interaction between hPDGFRβ and PfTRAP on the cell surface.

**Figure 3:**
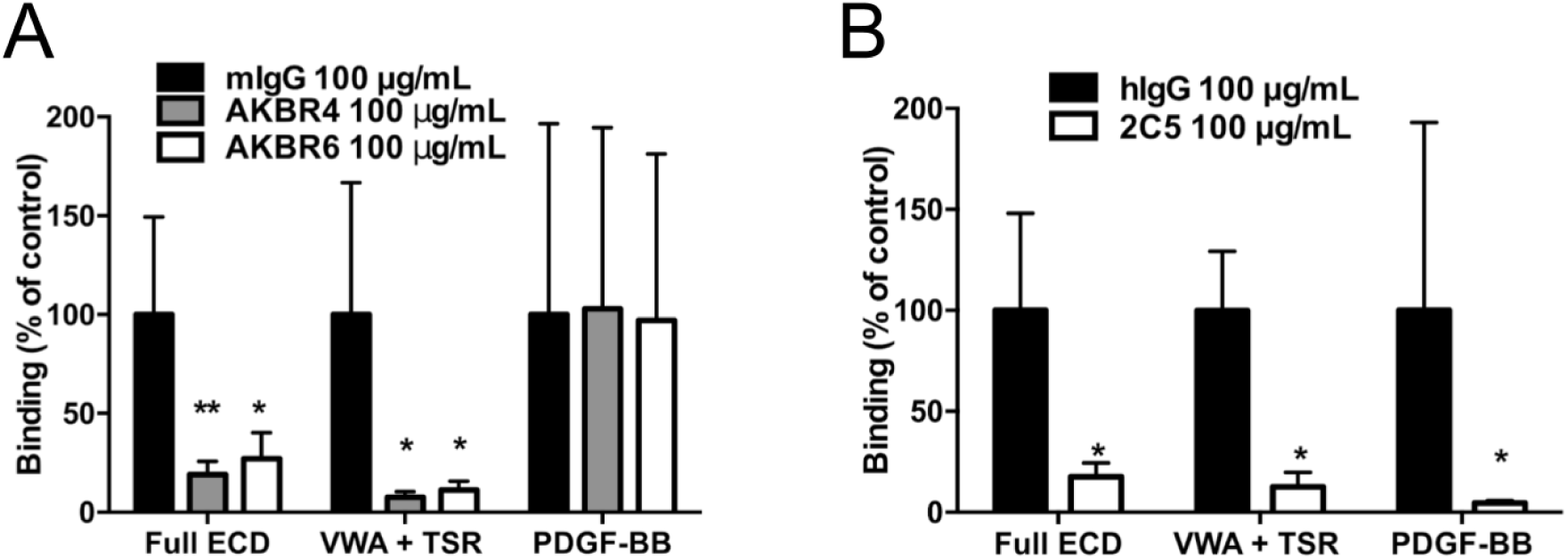
PfTRAP interaction with human PDGFRβ can be disrupted by parasite and host directed antibodies. (A) Anti-PfTRAP mAb AKBR4 or AKBR6 was incubated with 1000 nM PfTRAP full ectodomain (ECD) or VWA+TSR fragment, or 10 nM of the known PDGFRβ ligand hPDGF-BB for 30 minutes before addition to hPDGFRβ transfected cells in the flow assay. Binding of the Full ECD and VWA+TSR fragments was reduced by approximately 80% and 90%, respectively, while binding of human PDGF-BB was unaffected. (B) hPDGFRβ-transfected cells were incubated with anti-PDGFRβ mAb 2C5 for 30 minutes before addition of staining reagents as above with analysis by flow cytometry. Blocking hPDGFRβ inhibited binding of both PfTRAP fragments as well as the known ligand human PDGF-BB. Data are normalized to appropriate isotype controls and represent the mean ± SD from four independent experiments. Analysis by One-way ANOVA with Dunnet’s multiple comparisons test (A), and Mann Whitney test (B); *p<0.05, **p<0.01 compared to the isotype control.

Despite having shown binding of PfTRAP to cell-associated hPDGFRβ in our cell-based screen, we were unable to detect binding of PfTRAP to the recombinant soluble form of hPDGFRβ *in vitro* (Suppl. Fig. 6A), even when measured at high concentration of analyte or in the presence of excess divalent cations (Suppl. Fig. 6B).

In a recent report, the ectodomain of *P. falciparum* TRAP was shown to bind αv-integrin in a cell-based assay. The mechanism required divalent cations, presumably coordinated by the MIDAS motif of the TRAP VWA domain, and an integrin-binding RGD motif following the TSR domain (26). We therefore sought to distinguish αv-integrin-mediated interactions from those we found for hPDGFRβ. The HEK293F cells used in our assay endogenously express αv-integrin (Suppl. Fig. 7). Therefore, we measured PfTRAP binding to untransfected cells under conditions previously shown to inhibit binding to αv-integrin. PfTRAP ectodomain, indeed, bound to untransfected cells and this binding was disrupted in the presence of the divalent cation chelator EDTA as well as by anti-αv-integrin antibodies (Fig. 4A, B), in agreement with previous findings (26). Furthermore, our PfTRAP VWA+TSR fragment lacks the RGD-containing repeat region, and did not bind significantly to untransfected cells (Fig. 4B). This indicated that the binding to untransfected cells potentially was due to the interaction with the αv-integrin. We next tested the effect of EDTA and anti-αv-integrin antibodies on PfTRAP binding to cells transfected with hPDGFRβ. Both the full ectodomain and the VWA+TSR constructs of PfTRAP showed strong binding that was not affected by EDTA or anti-αv-integrin antibodies (Fig. 4C, D), indicating that the binding measured in hPDGFRβ-transfected cells was not due to binding endogenous αv-integrin, but was specific to hPDGFRβ.

**Figure 4:**
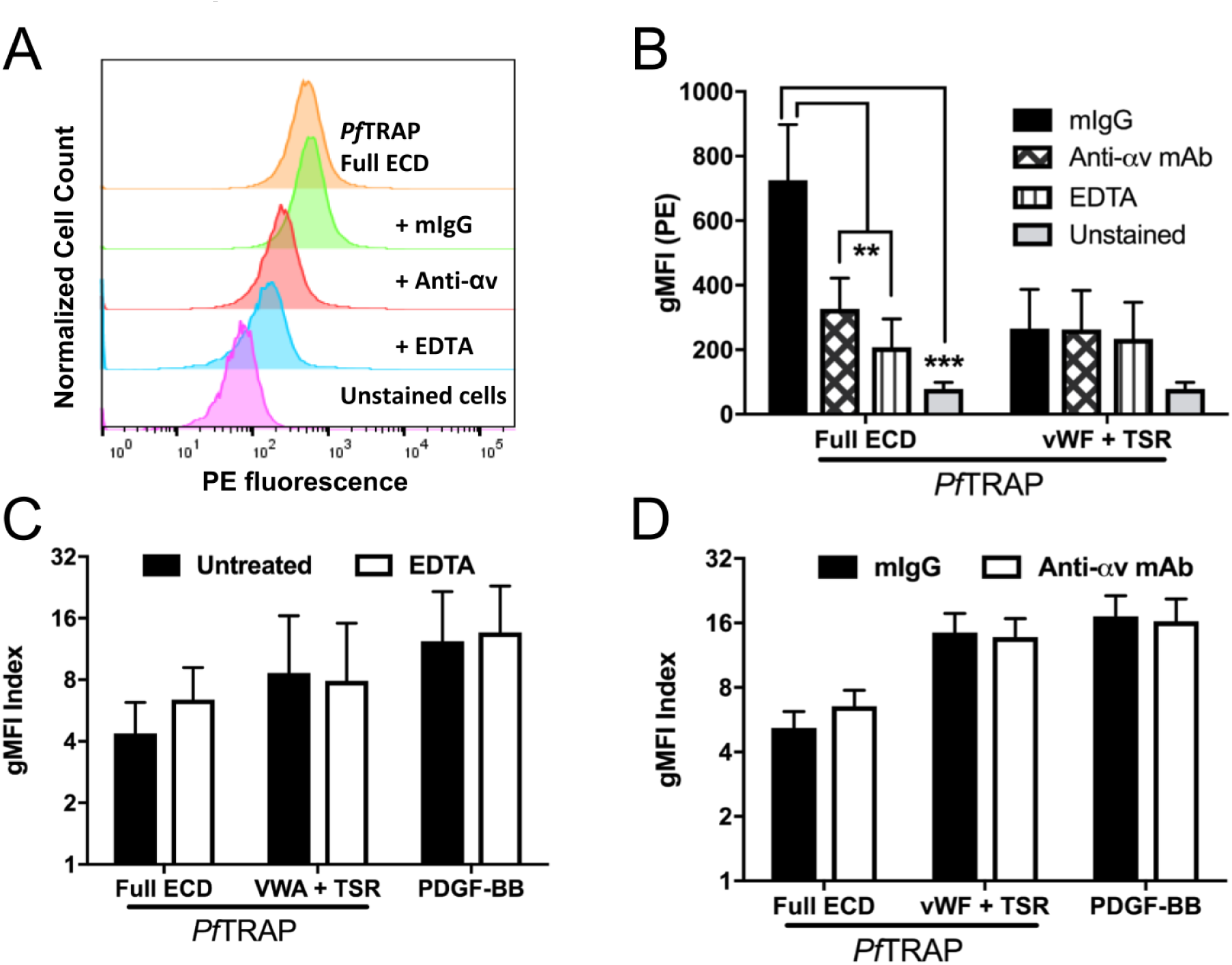
Divalent cations and αv-integrin do not contribute to PfTRAP binding of human PDGFRβ. (A, B) PfTRAP ectodomain (Full ECD) staining reagent at 1000 nM bound to untransfected HEK293F cells. This binding was inhibited by both 10 mM EDTA and 10 µg/mL of anti-αv-integrin mAb, but not an isotype control antibody. (B) The VWA+TSR fragment of PfTRAP did not bind significantly to untransfected cells and was not affected by EDTA or anti-αv-integrin antibodies. (C, D) Presence of 10 mM EDTA or 10 ug/mL anti-αv-integrin mAb did not significantly affect the binding of PfTRAP ECD (1000 nM), the VWA+TSR fragment (1000 nM) or the natural ligand PDGF-BB (10 nM) to HEK293F cells transfected with human PDGFRβ. Data are the mean ± SD from at least three independent experiments. Analysis by One-way ANOVA with Dunnet’s multiple comparisons test (B), and paired t-test (C-D); *p<0.05, **p<0.01, ***p<0.001.

Collectively these data show that optimal binding of PfTRAP to hPDGFRβ is mediated by the VWA and TSR domains acting together. We also show that this interaction can be disrupted using host- and parasite-directed antibodies and demonstrate that the binding of PfTRAP to hPDGFRβ-expressing cells is distinguishable from the recently discovered interaction with αv-integrins (26).

We next set out to test whether the TRAP:PDGFRβ interaction is important for sporozoite infection of the mammalian host. To this end, we tested passive transfer of mPDGFRβ blocking mAb 2C5 (which also blocks mouse mPDGFRβ) into mice, followed by sporozoite challenge via mosquito bite using the rodent malaria parasite *P. yoelii* (Suppl. Figs. 5C, 5D). Interestingly, mPDGFRβ blockade with mAb 2C5 did not have a significant effect on sporozoite infection of mice when compared to control. This result precipitated a further investigation into whether the TRAP:PDGFRβ interaction is conserved among *Plasmodium* species.

### Engagement of PDGFRβ is conserved across human-infective *Plasmodium* species but not in a rodent-infective *Plasmodium* species

We produced a recombinant ectodomain fragment for the human-infective malaria parasite *P. vivax* (PvTRAP), as well as full ECD and N-terminal adhesive domains (VWA+TSR) fragments for the rodent infective malaria parasite *P. yoelii* (PyTRAP) (Table 1; Suppl. Figs. 1C, 1D) and tested them for binding to cells overexpressing PDGFRβ of mouse (mPDGFRβ) and human (hPDGFRβ) origin. Binding of PvTRAP to hPDGFRβ, was seen in a concentration-dependent manner that was not significantly different from the binding of PfTRAP (Fig. 5A). We, however, observed no binding of either PyTRAP ectodomain or VWA+TSR to mPDGFRβ (Fig. 5B). This might explain why we observed that blocking the interaction in a rodent malaria infection model had no discernable effect on sporozoite infection (Suppl. Figs. 5C and 5D).

**Figure 5:**
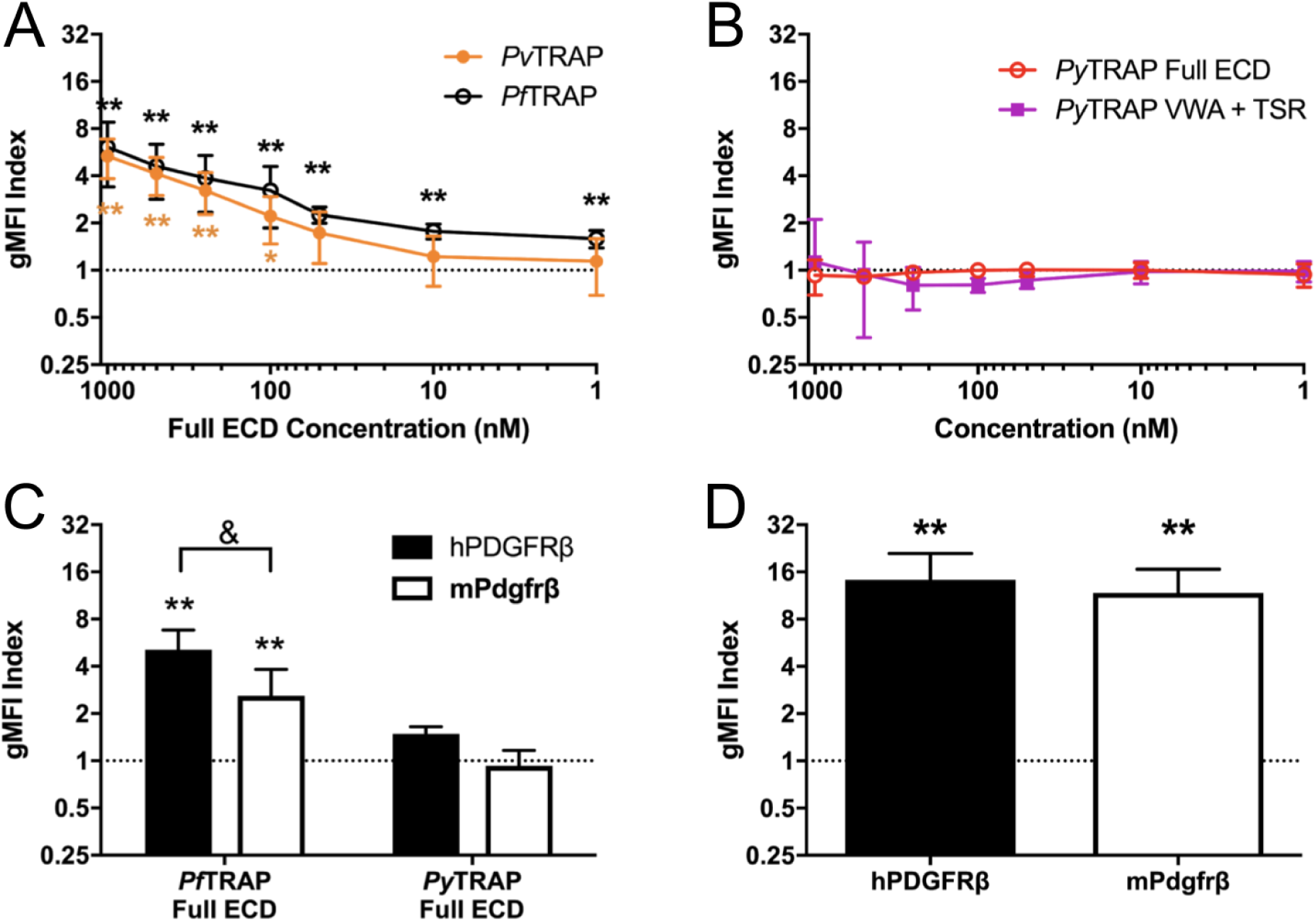
Engagement of PDGFRβ by TRAP distinguishes human and rodent infective *Plasmodium* species and is restricted at the level of TRAP. (A) PvTRAP ectodomain (ECD) staining reagent bound HEK293F cells transfected with human PDGFRβ (hPDGFRβ) in a concentration dependent manner that was not different from the binding of PfTRAP ECD. (B) PyTRAP ECD or a fragment containing the VWA+TSR domains did not bind to HEK293F cells transfected with mouse PDGFRβ (mPDGFRβ) at any concentration tested. (C) PfTRAP and PyTRAP ECD were tested at 1000 nM for binding to both hPDGFRβ and mPDGFRβ transfected HEK293F cells. PfTRAP ECD bound to both receptors, although less strongly to mPDGFRβ, while PyTRAP ECD did not bind to either receptor. (D) Known ligand for hPDGFRβ and mPDGFRβ, human PDGF-BB bound to both receptors at 10 nM. Data are the mean ± SD from at least three independent experiments. Asterisks indicate analysis by one sample t-test compared to the dashed line (gMFI index expected if no specific binding occurs), *p<0.05, **p<0.01; ampersand indicates analysis by t-test, ^&^p<0.05.

Finally, we assessed the cross-interaction of PyTRAP with hPDGFRβ, and PfTRAP with mPDGFRβ. The PfTRAP fragment bound significantly to mPDGFRβ, albeit only at approximately 50% of the level of binding to hPDGFRβ (Fig. 5C). No binding of PyTRAP to either mPDGFRβ or hPDGFRβ was observed. Both human and mouse PDGFRβ bound the human PDGF-BB ligand to a similar extent (Fig. 5D), ruling out gross discrepancies in protein localization or expression levels affecting these results.

## Discussion

Over the last decade, numerous *Plasmodium* proteins have been identified that play a role in the sporozoites’ journey from the site of deposition in the skin to their destination in the liver and their ultimate infection of host hepatocytes (27). Furthermore, host hepatocyte surface proteins CD81, SR-B1 and EphA2 have been described as critical factors for infection of hepatocytes (28–30). Yet, linking host receptors to sporozoite ligands has been exceedingly difficult, and sporozoite protein/host cell protein interaction pairs that might play a role in host infection remain virtually unknown. We have herein identified PDGFRβ as a novel host receptor for the essential sporozoite adhesin TRAP by screening >4300 human plasma membrane proteins. The PDGFRβ:TRAP interaction can be blocked with mAbs to either the parasite protein or the host receptor, requires both TRAP VWA and TSR domains for efficient binding, and is independent of the TRAP MIDAS and RGD motifs. Intriguingly, the PDGFRβ interaction with TRAP is conserved among the major human-infective *P. falciparum* and *P. vivax* parasites but not for the rodent-infective parasite *P. yoelii*.

Binding of recombinant PfTRAP to hPDGFRβ was detectable in our cell-based assay in a concentration-dependent manner. However, we were unable to detect binding of recombinant TRAP to the purified soluble form of PDFGRβ, despite detecting a robust interaction with its known ligand PDGF-BB. It is possible that the conformation of the membrane-anchored PDGFRβ differs from that of its soluble counterpart, or that TRAP interaction requires presence of additional proteins associated with PDGFRβ.

Our findings complement the recent study by Dundas and colleagues (26) in which αvβ3 integrin was identified as a host receptor for TRAP. It is likely that this interaction was not detected in our human surface protein screen because only individual proteins were overexpressed. We have independently validated the binding of PfTRAP to αv-containing integrins, which have basal expression in our cell-based system. However, we have shown that binding to hPDGFRβ was distinct as indicated by our findings that (i) binding to hPDGFRβ did not require the RGD motif containing repeat region, (ii) binding was insensitive to anti-αv antibodies and (iii) unlike the interaction with αv-containing integrins (26), binding did not require divalent cations. Furthermore, the interaction with hPDGFRβ was conserved for TRAP of *P. falciparum* and *P. vivax*. In contrast, the interaction between integrin αvβ3 and the TRAP of *P. vivax* had not been observed by Dundas *et al*. (26). Thus, it is highly likely that the interactions of TRAP with αvβ3 and PDGFRβ are distinct and might serve different biological functions during the infection process.

In order to directly establish the potential importance of the TRAP:PDGFRβ interaction, we initially attempted to block the interaction *in vivo* and measure the effect on sporozoite transmission success, using the *P. yoelii* mouse malaria model. Interestingly, however, *P. yoelii* TRAP failed to show significant binding to hPDGFRβ or mPDGFRβ (Fig. 5C). Therefore, it is probable that the interaction between TRAP and PDGFRβ is not conserved in *P. yoelii*, and this will not be an adequate model to study the biology of TRAP:PDGFRβ interaction. Thus, our findings are adding to the emerging picture of differences in the conservation of host receptor engagement by orthologous effector proteins between human-infective and rodent-infective *Plasmodium* species (26, 31).

PDGFRβ plays a major role in angiogenesis and maintenance of vascular integrity, and was shown to be expressed by lymphatic endothelial cells as well as pericytes supporting endothelial cells during development (32–34). Additionally, PDGF receptors have long been recognized to play an important role in cutaneous wound healing (35). Consistent with these published vascular expression patterns, we detected PDGFRβ expression in a perivascular distribution in human skin. Expression was strong in pericytes that undergird the vascular endothelium, which in turn was unequivocally identifiable with the endothelial cell marker CD31. Likewise, liver PDGFRβ expression was restricted to vasculature. Thus, it is possible that the TRAP:PDGFRβ interaction may have a role in mediating sporozoite recognition of the vasculature. However, our findings indicate that TRAP:PDGFRβ interaction is not conserved among the different *Plasmodium* species, and that further functional studies of this interaction must be undertaken in model systems that recapitulate the human vascular compartment.

Combined with recent studies, our findings provide a more complete understanding of the potential molecular interactions of TRAP with host receptors. While the biological consequences of the TRAP:PDGFRβ interaction remain to be fully deciphered, our work provides a novel molecular interaction lead that could mediate the initial infection of the mammalian host.

## Experimental procedures

### Recombinant Protein Production

Production of recombinant proteins in suspension HEK293F cells cultures was performed, as previously described (28, 36, 37). Briefly, each expression construct contained codon-optimized sequences encoding a tissue plasminogen activator signal peptide (38), the full extracellular domain or subdomains from *P. yoelii* TRAP (PyTRAP) (PlasmoDB:PYYM_1351500), *P. falciparum* TRAP (PfTRAP) (PlasmoDB: PF3D7_1335900) or *P. vivax* TRAP (PvTRAP) (Table 1), and a C-terminal His_8_ tag followed by an AviTag (39) cloned into the pcDNA-3.4 vector backbone (Thermo Fisher, Waltham, MA, USA). Constructs encoding tagged human PDGF-B (residues 82–190) (purified as homodimeric hPGDF-BB), and the tagged ectodomain of hPDGFRβ (residues 33–532) were similarly created.

The PvTRAP sequence was based on that observed by proteomic analysis of a Thai *P. vivax* salivary gland sporozoite sample bearing the VK247 haplotype of CSP (40). Protein sequence polymorphisms in the field isolate were identified by augmenting the *P. vivax* Sal-1 genome (41) used for the protein sequence database search with the *P. vivax* P01 genome (42) as well as sequence polymorphisms observed in RNA-seq analyses of 19 different Thai field isolates (43). Additionally, S→A and T→A substitutions were made in NX[S/T] motifs to prevent N-linked glycosylation, as this modification has not been shown to occur in *Plasmodium* (44, 45) and is absent from TRAP *in vivo* (46).

Constructs were introduced into HEK293F cells by high-density transfection with PEI MAX 40K (Polysciences, Inc., Warrington, PA, USA) (47, 48), and culture supernatants containing the recombinant proteins were harvested after 5 days. Proteins were purified by Ni^2+^-affinity chromatography followed by size-exclusion chromatography. Subsequently, a BirA biotin-protein ligase kit (Avidity, LLC; Aurora, CO, USA) was used to biotinylate protein samples, as previously described (37). Biotinylated proteins were re-purified by size-exclusion chromatography to remove free biotin, and multimerized using streptavidin-phycoerythrin (ProZyme) by mixing 1 molar-equivalent of streptavidin-phycoerythrin with 4 molar-equivalents of biotinylated protein.

### Retrogenix® Cell Microarray Screen

Screening for PfTRAP binding proteins was performed using the Retrogenix Cell Microarray technology. Initially 4336 expression vectors, each encoding a unique human plasma membrane protein and the ZsGreen1 fluorescent protein, were pre-spotted on glass slides. Human HEK293 cells were grown over the arrays and reverse-transfected, resulting in the over-expression of each membrane protein and ZsGreen1. Slides were then incubated with QDot ITK Carboxyl Quantum Dots, 585 nM (Life Technologies cat # Q21311MP) which were pre-coupled with Ni-NTA and coated with His-tagged PfTRAP, giving a final PfTRAP concentration of 100 nM. PfTRAP binding proteins were identified by fluorescence imaging (at ∼500 nm wavelength for ZsGreen1 and at ∼585 nm for QDots) analysed using ImageQuant software (GE). For confirmation and specificity testing, expression vectors encoding the PfTRAP binding proteins were re-spotted on new slides, and HEK293 cells were reverse-transfected. Slides were then treated with the same Ni-NTA-coupled QDots coated with His-tagged PfTRAP as before, or with His-tagged PDL1 (each at 100 nM final concentration), or with no ligand conjugated. Interactions were analysed by fluorescence imaging.

### Protein binding measurements

Protein interactions were analyzed using biolayer interferometry on an Octet QK^e^ instrument (ForteBio). Streptavidin (SA) biosensors were used to immobilize biotinylated recombinant ligand molecules followed by equilibration in 10x Kinetics Buffer (PBS + 0.1% BSA, 0.02% Tween-20 and 0.05% sodium azide), detection of association of recombinant analyte proteins diluted in the same buffer, and detection of dissociation in analyte-free 10x Kinetics Buffer.

Experiments requiring the use of salts of divalent metals required phosphate to be substituted with HEPES (since phosphate formed poorly soluble salts with several of the metal ions tested). In those cases, we used HBS-P (20 mM HEPES, pH 7, 150 mM NaCl, 0.1% BSA, 0.02% Tween-20) as the base buffer.

### Flow Cytometry Binding Assays

To further analyze TRAP binding, biotinylated recombinant TRAP proteins (Table 1) were oligomerized around a streptavidin-conjugated Phycoerythrin (PE) (PJRS2, Prozyme) creating TRAP-PE staining complexes. Sequence verified human (clone IMAGE:5309813) and mouse (clone IMAGE:30060666) cDNAs for PDGFRβ were obtained from the Dharmacon Mammalian Gene Collection (GE Healthcare) for creation of surface-expressed GFP fusion receptor constructs. The open reading frame, including the transmembrane region and a short intracellular linker, were cloned into a pcDNA3.4 expression vector (Thermo Fisher) in which the intracellular tyrosine kinase domain of the receptor was replaced with Cycle3 GFP. Approximately 3 × 10^6^ HEK293F cells were transfected with one of these constructs using 293fectin Transfection Reagent (Thermo Fisher) or 293-Free (EMD Millipore), according to the manufacturer’s instructions (Thermo Fisher) and maintained in FreeStyle 293 serum-free medium (Thermo Fisher). Two days after transfection approximately 5 × 10^5^ cells were transferred to 96-well plates, pelleted at 500 × g for 3 minutes, then resuspended and incubated with various concentrations of protein complexes in FACS buffer (PBS + 2% FBS) for 30 minutes on ice. Cells were then washed three times with FACS buffer before analysis without fixation on an LSR II Flow Cytometer (Becton Dickinson) using FlowJo (TreeStar). Specificity for binding to the transfected receptor was measured by the geometric mean fluorescent intensity (gMFI) of PE on transfected (GFP^+^) cells normalized to the untransfected (GFP^−^) cells in the same well, termed the gMFI Index. Experiments with the known ligand for PDGFRβ, a dimer of the human Platelet-Derived Growth Factor-B (hPDGF-BB), and mock transfected cells were included as controls in the assays. Experiments testing the role of divalent metals in binding were done in presence of 10 mM EDTA. Blocking of the integrin αv binding was performed using the mouse monoclonal 272-17E6 antibody (Abcam).

### Recombinant antibody production

For PfTRAP blocking experiments monoclonal antibodies recognizing PfTRAP were produced as previously described (37). Briefly, BALB/cJ mice were immunized intramuscularly three times, three weeks apart, with 20 µg of PfTRAP full ectodomain formulated in Adjuplex (Sigma Aldrich). Two weeks following the final immunization, spleens from PfTRAP immunized mice were harvested, dissociated, and antigen-specific B cells sorted by FACS into 96-well plates for subsequent culture. Sorted cells were stimulated to secrete antibody and supernatants screened for antigen-binding antibodies by ELISA. Antibody sequences encoding the variable regions of heavy and light chains were rescued from the positive wells using RT-PCR and fused to constant-region encoding DNA fragments to build expression constructs.

For PDGFRβ receptor blocking experiments, a previously published monoclonal antibody identified from a phage display library (clone 2C5) and capable of neutralizing both mouse and human PDGFRβ (25) was expressed as a recombinant human IgG1. Briefly, the variable heavy- and light-chain sequences were obtained from Creative Biolabs (NY, USA) and cloned directly into antibody expression vectors containing constant-region sequences.

Antibodies were expressed using the HEK293F culture, as described above for protein production. Following 5 days in culture, supernatants were cleared by centrifugation, supplemented with NaCl (+350 mM, final concentration), and applied over Protein G Sepharose Fast Flow (GE Lifesciences). Protein was eluted using 0.1 M glycine (pH 2.7), and the pH was immediately neutralized using 1 M dibasic sodium phosphate.

### Immunohistochemistry

Localization of PDGFRβ in formalin-fixed paraffin-embedded normal human skin and liver sections (Amsbio, MA, USA) was performed using a EnVision G2 Doublestain System, Rabbit/Mouse (Agilent). This staining procedure can be used for sequential double staining of two different antigens with primary antibodies raised in the same species by incorporating a blocking step between the first and second antibody incubations. Briefly, sections were deparaffinized and rehydrated by standard methods, antigens retrieved by immersion in boiling 10 mM sodium citrate (pH 6.0) for 10 minutes, and endogenous peroxidase activity quenched according to the manufacturer’s instructions. Sections were then blocked and permeabilized in Protein Block Serum-Free (Agilent) for 20 min at room temperature before incubation with primary rabbit monoclonal antibody against PDGFRβ (Abcam; clone Y92). Sections were washed twice in TBS-Tween 0.05% (20 mM Tris pH 8, 150 mM NaCl, 0.05% Tween-20) for 5 min. Polymer/HRP (Agilent) was added to the sections and sections were incubated for 30 min at room temperature in the dark. Sections were washed twice with TBS-Tween 0.05% for 5 min each. DAB chromogen (diluted 1 drop every 1 mL DAB buffer) was added to the sections, and slides were incubated in the dark for 20 min. Sections were washed with distilled water to stop the reaction. For staining of vasculature, sections were stained with the rabbit monoclonal antibody against the endothelial marker CD31 (Abcam; clone EPR3094). Sections were washed with TBS-Tween 0.05% for 5 min and rabbit/mouse link (Agilent) was added to slides and incubated for 15 min at room temperature in the dark. Sections were washed twice in TBS-Tween 0.05% for 5 min and polymer-AP was added. Sections were incubated for 30 mins at room temperature in the dark followed by two washes with TBS-Tween 0.05% 5 min. To visualize CD31, Vector Blue substrate (Vector Laboratories) was prepared according to manufacturer’s instructions and added to the sections for 20 mins. Reaction was stopped with water. PDGFRβ straining (brown) and CD31 staining (blue) were visualized and imaged using the Keyence BZ-X710 microscope.

### Mosquito Rearing and Sporozoite Production

*Anopheles stephensi* mosquitoes were reared according to standard protocols and adult females were infected with *Plasmodium* parasites 3-7 days after their emergence. For *P. yoelii* sporozoite production, female 6-8-week-old Swiss Webster mice were injected with blood stage 17XNL parasites constitutively expressing GFP/Luciferase (*Py*GLuc) (49) to begin the growth cycle, and infected mice were used to feed adult female *Anopheles stephensi* mosquitoes after gametocyte exflagellation was observed. Blood-fed mosquitoes were then maintained according to standard protocols before their use in animal challenge studies.

### mAb 2C5 passive transfer protection studies

All experiments involving animals were performed under protocols approved by the Institutional Animal Care and Use Committee at the Seattle Children’s Research Institute. Mice received two, 800-µg injections of mAb 2C5 or isotype control human IgG before infectious mosquito bite challenge. The first injection was i.p. 18 hours before challenge to allow distribution of the antibody throughout the tissues, while the second was i.v. 1 hour before to ensure a high serum antibody concentration at the time of challenge. Host directed antibodies against CD81 have been shown to prevent malaria infection with a single dose of 800 µg the day before infection (50). Mouse infection and analysis of malaria liver stage burden was as previously described (49, 51). Briefly, BALB/cJ mice were anesthetized and infected with the bites of 20 *Py*GLuc infected mosquitoes, respectively. Malaria liver stage burden was measured at the end of liver stage development (45 hours post infection) using an In Vivo Imaging System (IVIS) to determine if blocking PDGFRβ resulted in lower malaria liver stage infection. From day 3 post infection, mice were monitored daily for the presence of erythrocytic stages by Giemsa staining of thin blood smears.

### Statistical Analysis

All statistical analyses were performed using GraphPad Prism software (version 6), and differences considered statistically significant when p was <0.05.

## Data availability

All data presented are contained within the article and the Supporting Information.

## Acknowledgements

We would like to thank the Seattle Children’s Research Institute insectary staff, principally Heather Kain, for the help with mosquito production. We also thank Drs. S. Mikolajczak and E. Flannery (Novartis Institute for Tropical Diseases) for assistance with the design of the *P. vivax* TRAP protein.

## Funding and additional information

Our work was supported by National Institutes of Health Grants R01AI134956 (S.H.I.K), R01AI117234 (D.N.S. and S.H.I.K), K25AI119229 (K.E.S), R01GM087221 (R.L.M.). The content is solely the responsibility of the authors and does not necessarily represent the official views of the National Institutes of Health.

## Footnotes

## The abbreviations used are

TRAP: thrombospondin-related anonymous protein;
VWA: von Willebrand factor A;
MIDAS: metal ion-dependent adhesion site;
TSR: thrombospondin repeat;
HSPGs: heparan sulfate proteoglycans;
hPDGFRβ: human platelet-derived growth factor receptor β;
gMFI: geometric mean fluorescence intensity.

## Supporting information

**Supplementary Figure 1:**
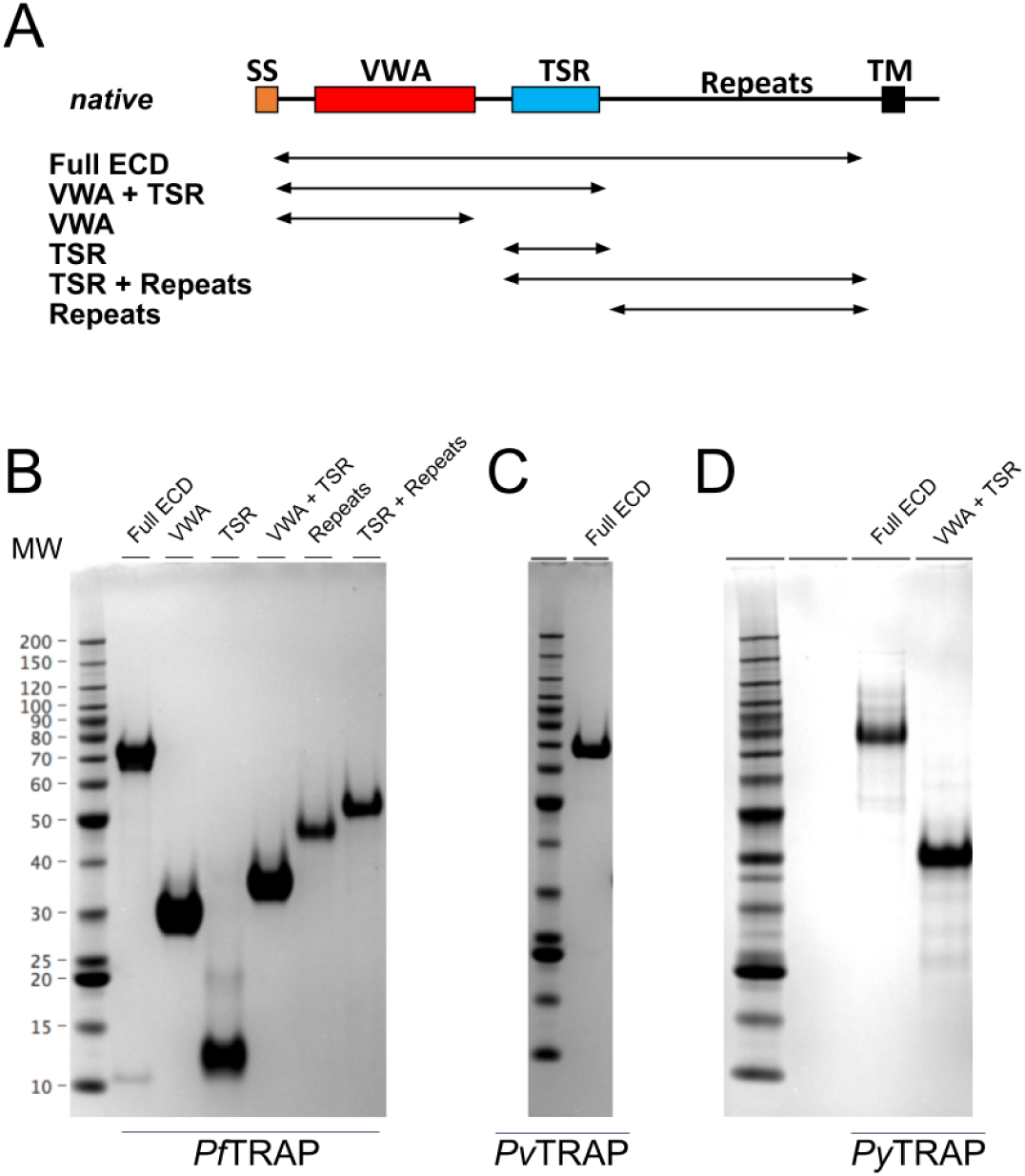
Characterization of recombinant *Plasmodium* TRAP ectodomains fragments. (A) Schematic representation of deletion constructs spanning the ectodomain of TRAP. SDS-PAGE of the six constructs produced using the *P. falciparum* (B), *P. vivax* (C), and *P. yoelii* (D) sequences. Labels for the molecular weight standards are shared across panels B–D.

**Supplementary Figure 2:**
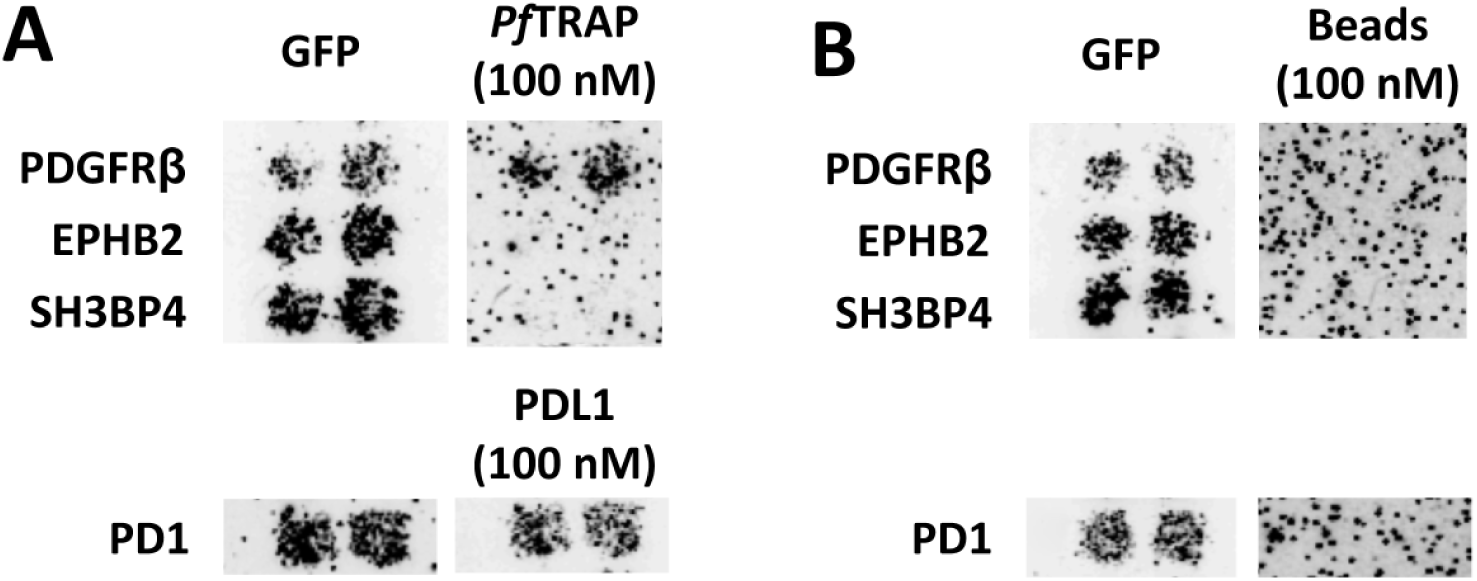
*Pf*TRAP binds to PDGFRβ expressed in HEK293 cells using the Retrogenix® Cell Microarray platform. Hits from primary binding screens were re-arrayed for reverse transfection into HEK293 cells in duplicate for secondary specificity and reproducibility screens in conjunction with a GFP reporter construct for successful transfection. Fluorescent beads conjugated with PfTRAP full ECD identified specific binding to PDGFRβ transfected cells (A) that was not observed with unconjugated beads (B). Positive control ligand, PDL1, oligomerized to fluorescent beads bound strongly and specifically to its known interaction partner PD1.

**Supplementary Figure 3:**
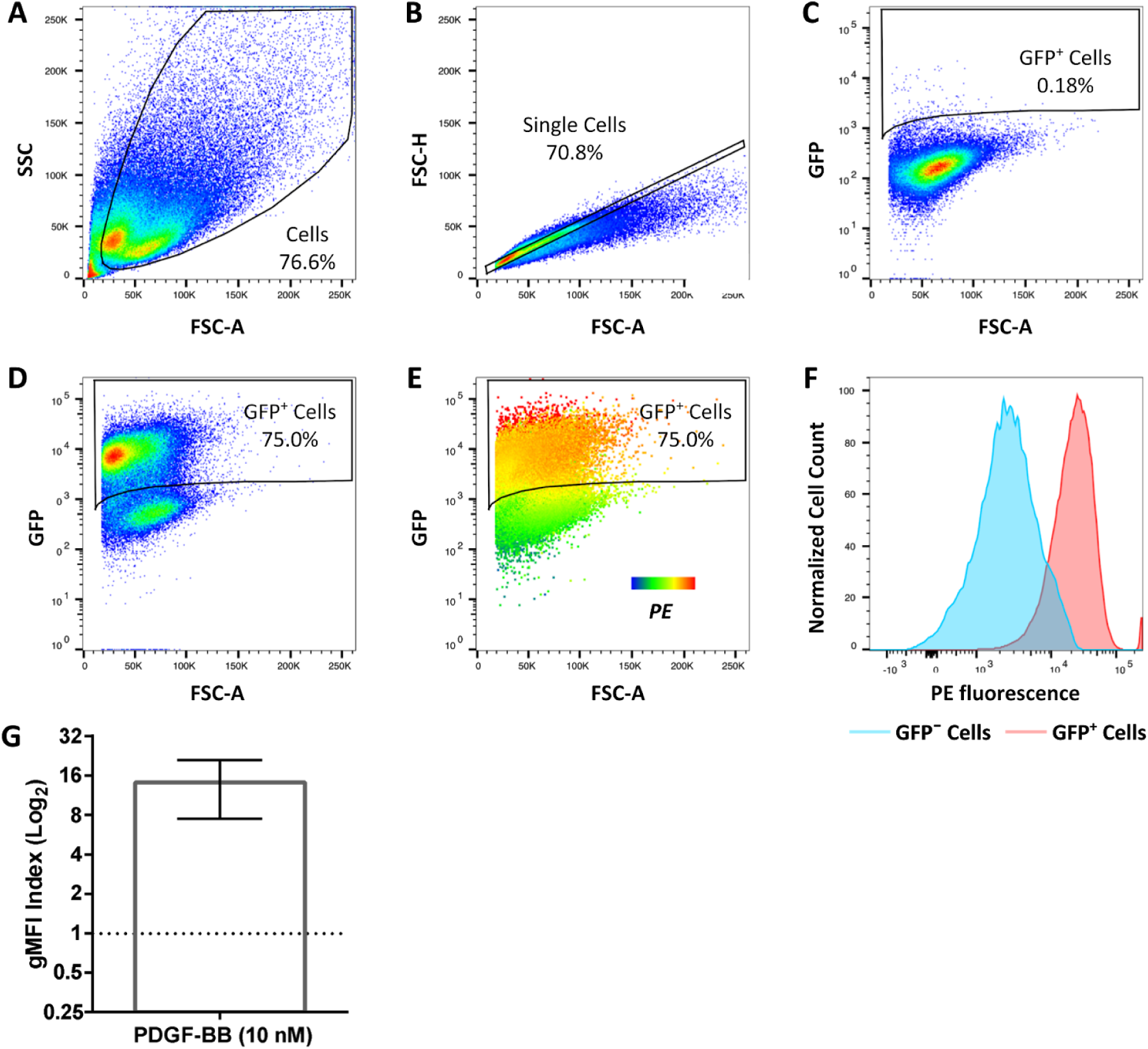
Analysis of ligand binding by flow cytometry. Biotinylated ligand (PDGF-BB in this example) was oligomerized around a streptavidin-conjugated phycoerythrin (PE) staining reagent and incubated with HEK293F cells for 30 minutes before analysis by flow cytometry. Cells were first separated from debris (A) and single cells selected for analysis (B). An untransfected cell sample was used to set the GFP^+^ transfected cell gate (C) from which cells transfected with a GFP-fusion of PDGFRβ and untransfected GFP^−^ cells from within the same sample were identified (D). The geometric mean fluorescent intensity (gMFI) of PE staining was used to determine the binding of the oligomerized staining reagent to the transfected and untransfected cell population, for which the positive control ligand human PDGF-BB bound strongly and preferentially to the PDGFRβ transfected cell population (E). Specificity for binding to the transfected receptor population was measured as the ratio of the gMFIs from GFP^+^ and GFP^−^ cells reported as the ‘gMFI Index’, and measurements were taken over multiple experiments to ensure the reproducibility of the results (F–G). Data in A–F show a representative experiment. Data in G are from 7 independent experiments and represent the mean ± SD; the dashed line represents the gMFI index expected if no specific binding occurs.

**Supplementary Figure 4:**
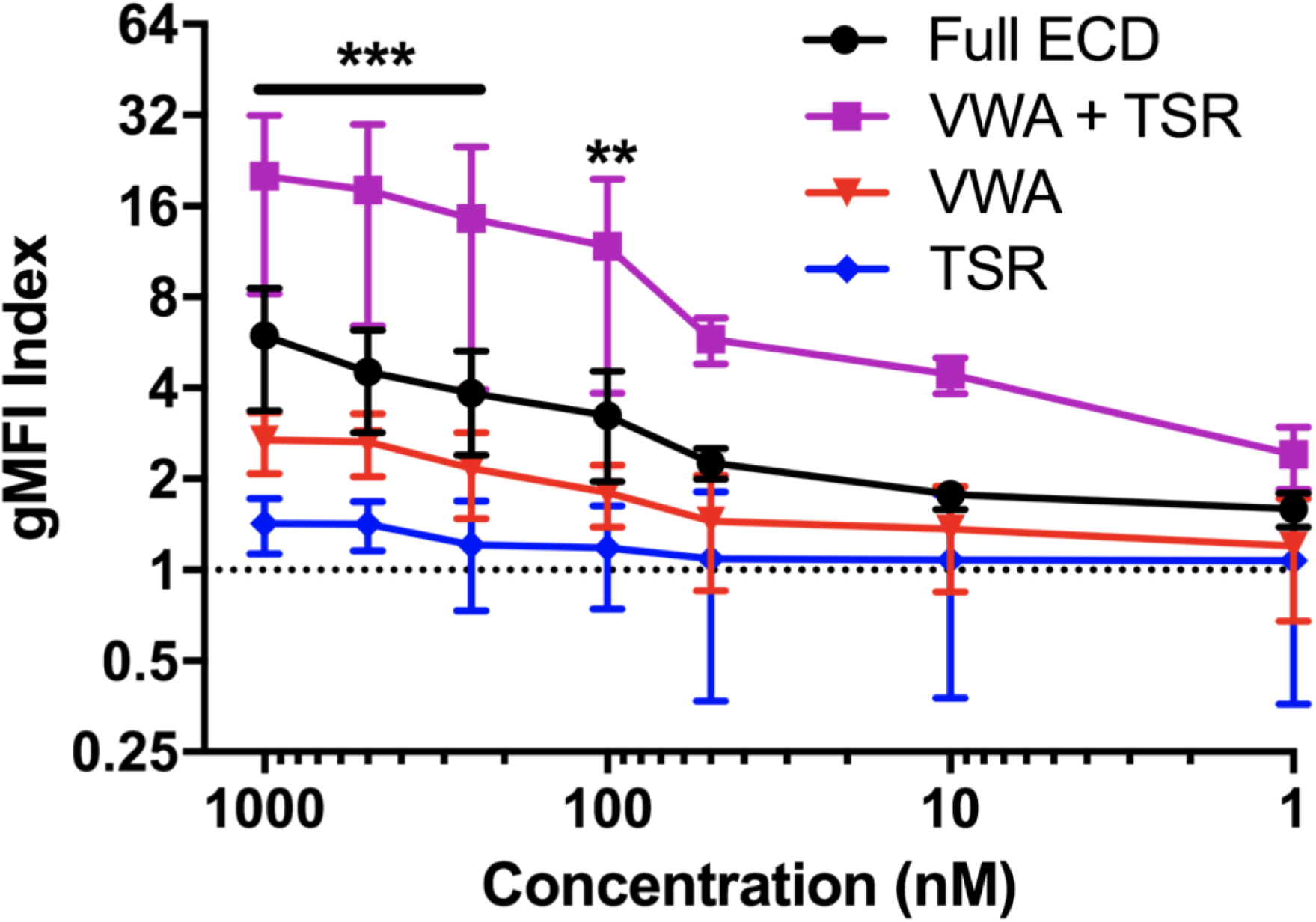
Titration of PfTRAP fragment binding to hPDGFRβ in the flow cytometry binding assay. The gMFI index of VWA+TSR fragment bound to PDGFRβ transfected cells was greater than all other fragments at concentrations at or above 100 nM. Data are the same as those in Figure 1 and represent the mean ± SD from at least five independent experiments. Analysis by Two-Way ANOVA with Bonferroni’s multiple comparisons test; asterisks indicate the VWA+TSR fragment is different from all others, **p<0.01, ***p<0.001.

**Supplementary Figure 5:**
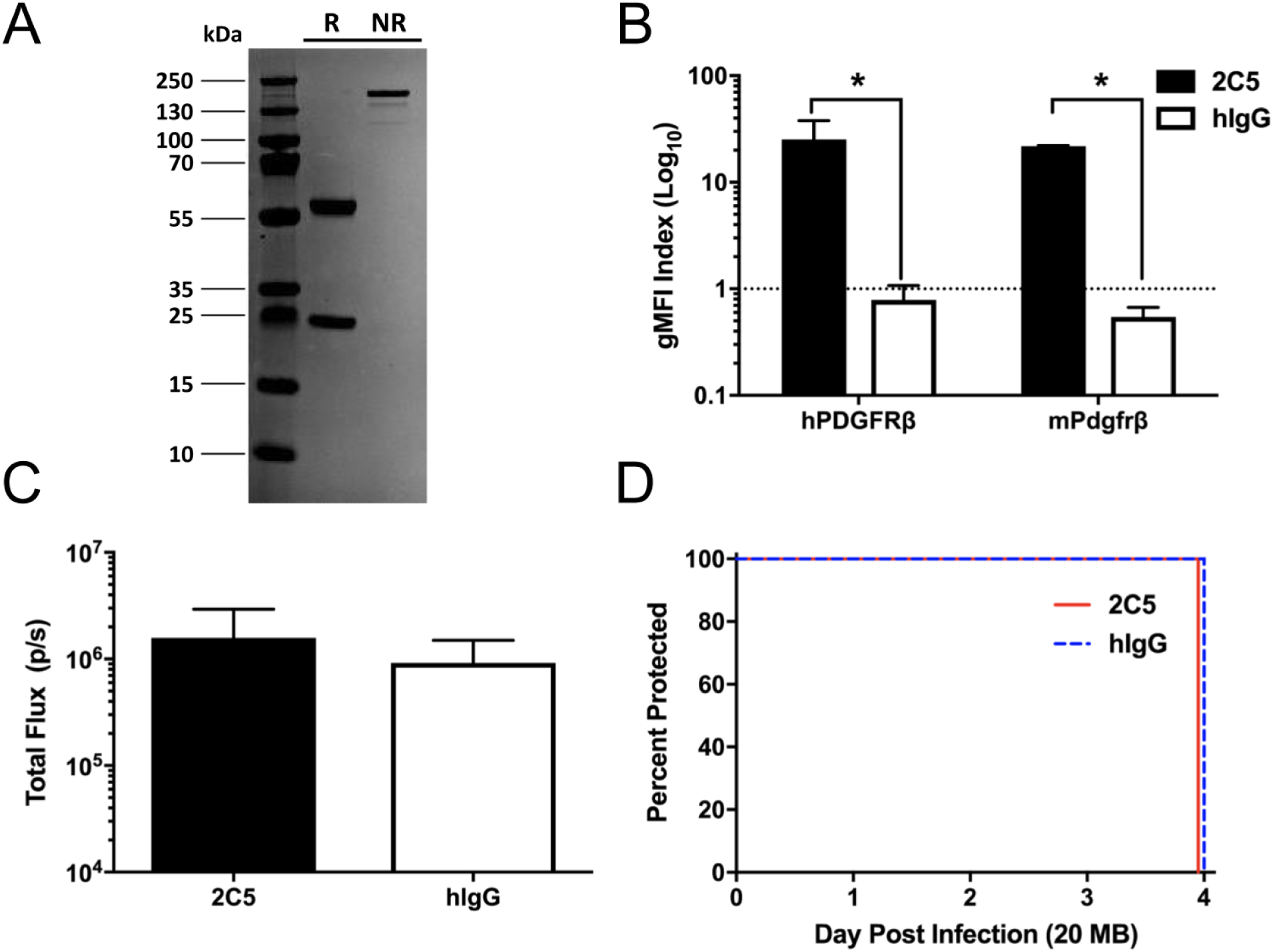
Validation of anti-PDGFRβ mAb 2C5 and its use in *P. yoelii* passive transfer studies. (A) mAb 2C5 was produced as a human IgG1 and analyzed by SDS-PAGE under reducing (R) and non-reducing (NR) conditions. (B) Cells were transfected with human or mouse PDGFRβ (hPDGFRβ and mPDGFRβ, respectively), then incubated with 100 μg/mL mAb 2C5 or isotype control before washing and staining with an anti-human antibody to measure mAb 2C5 specificity for PDGFRβ. Data are the mean ± SD from at least 2 independent experiments with analysis by t-test; *p<0.05. (C-D) Eighteen hours and one hour before challenge with 20 bites from *P. yoelii* infected mosquitoes, BALB/cJ mice received 800 μg mAb 2C5 (n=5) or isotype control (n=5; 1.6 mg of antibody total). There was no difference in the malaria liver stage burden at 45 hours post infection or delay to blood stage patency.

**Supplementary Figure 6:**
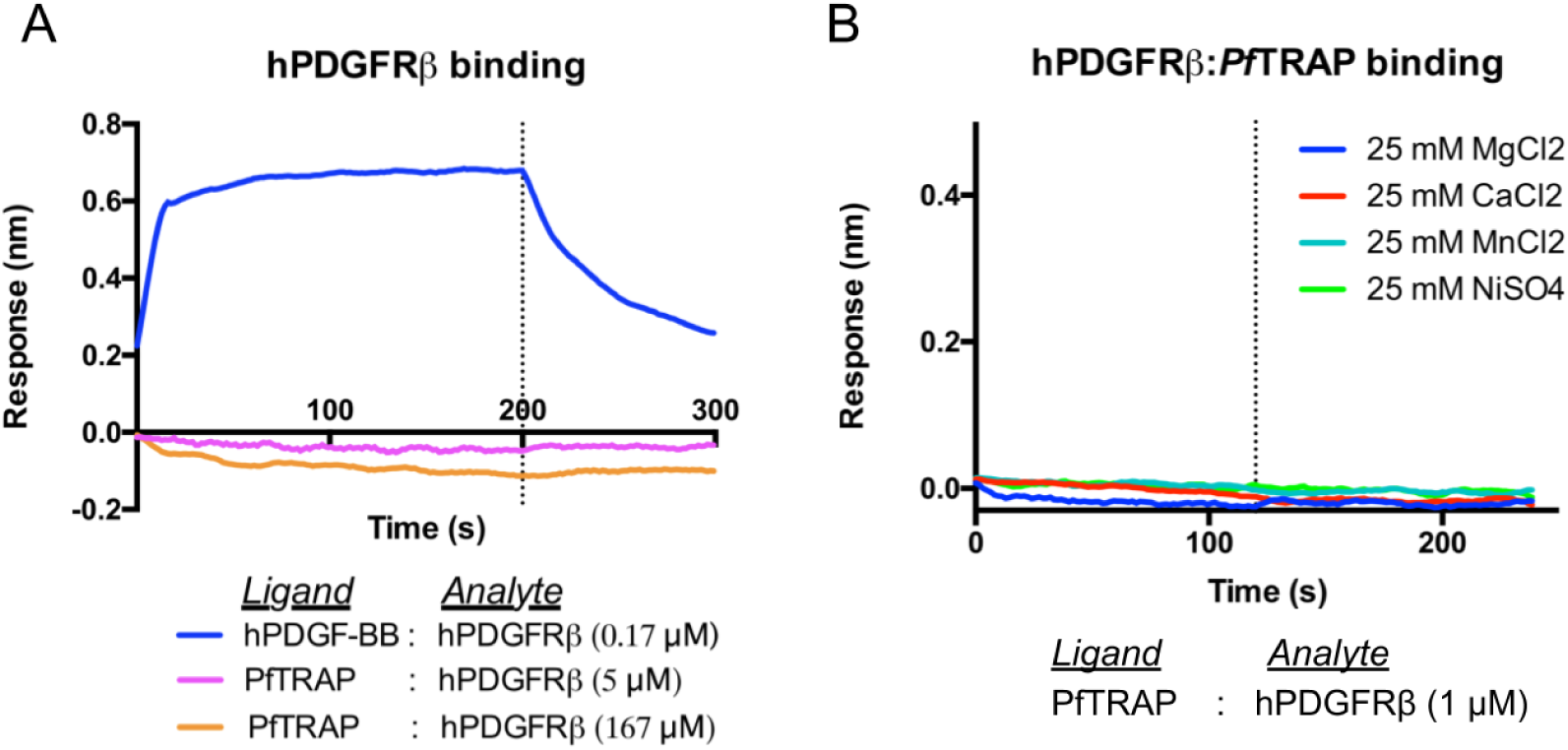
Purified recombinant human PDGFRβ does not bind purified recombinant PfTRAP in biolayer interferometry experiments. (A) Solutions containing recombinant biotinylated PfTRAP or hPDGF-BB (ligands) were used to derivatize streptavidin biosensors prior to incubating them in solutions containing recombinant hPDGFRβ (analyte) at the indicated concentrations for 200s, subsequently incubating them in reference buffer for another 100s. (B) Experiments using biotinylated PfTRAP as ligand were performed in the presence of 25-mM concentrations of divalent cations, as indicated in the legend. Sensorgrams in each experiment were reference-subtracted using data from an analyte-free reference well.

**Supplementary Figure 7:**
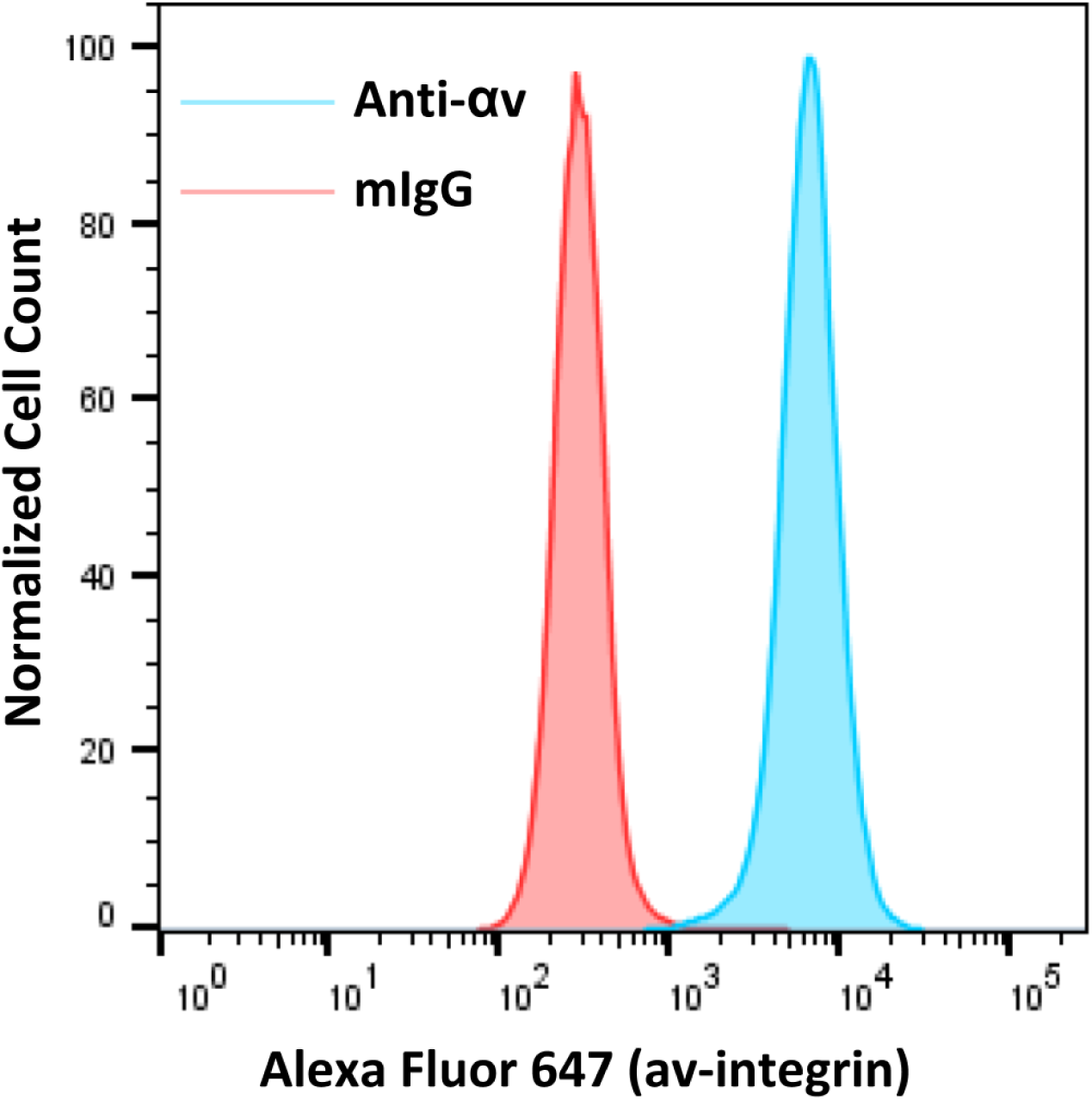
HEK293F cells express αv-integrin. Untransfected HEK293F cells were incubated with 10 µg/mL anti-αv-integrin antibody or mouse IgG (mIgG) before washing and staining with an anti-mouse Alexa Fluor 647 antibody. Cells incubated with the anti-αv-integrin antibody were significantly brighter than the isotype control.

